# Dissecting the chain of information processing and its interplay with neurochemicals and fluid intelligence across development

**DOI:** 10.1101/2022.11.04.515156

**Authors:** George Zacharopoulos, Francesco Sella, Uzay Emir, Roi Cohen Kadosh

**Affiliations:** Wellcome Centre for Integrative Neuroimaging, Department of Experimental Psychology, University of Oxford, UK; School of Psychology, Swansea University, UK; Centre for Mathematical Cognition, Loughborough University, UK; School of Health Sciences, College of Health and Human Sciences, Purdue University, USA; School of Psychology, University of Surrey, Guildford, UK

**Keywords:** visuomotor processing, fluid intelligence, glutamate, GABA, development, frontoparietal network

## Abstract

Previous research has highlighted the role of glutamate and gamma-aminobutyric acid (GABA) in perceptual, cognitive, and motor tasks. However, the exact involvement of these neurochemical mechanisms in the chain of information processing, and across human development, are unclear. In a cross-sectional longitudinal design, we used a computational approach to dissociate cognitive, decision, and visuomotor processing in 293 individuals spanning early childhood to adulthood. We found that glutamate and GABA within the intraparietal sulcus (IPS) explained unique variance in visuomotor processing, with higher glutamate predicting poorer visuomotor processing in younger participants but better visuomotor processing in mature participants, while GABA showed the opposite pattern. These findings were functionally, neuroanatomically, and neurochemically specific, and were replicated ~1.5 years later and were generalized in two further different behavioral tasks. Using resting fMRI, we revealed that the relationship between IPS neurochemicals and visuomotor processing is mediated by functional connectivity in the visuomotor network. We then extended our findings to high-level cognitive behavior by predicting fluid intelligence performance. We showed that fluid intelligence performance is explained by IPS GABA and glutamate and is mediated by visuomotor processing, and moderated by the developmental stage. These results provide an integrative biological and psychological mechanistic explanation that links cognitive processes and neurotransmitters across human development and establishes their involvement in intelligent behavior.

## 1. Introduction

Previous research has highlighted the role of glutamate and gamma-aminobutyric acid (GABA) in the function and malfunction of the central nervous system underpinning cognitive, decision, and visuomotor processing (1–6). However, little is known about the neurochemical mechanisms shaping these processes across development in humans. An influential computational framework has been extensively applied in recent years as it allows the behavioral dissection between cognitive, decision, and visuomotor processes. This framework is based on the drift–diffusion model and analyzes reaction times and accuracy, usually in two-choice response time tasks (7–9).

The diffusion parameters, which we describe in detail in the following paragraph, have shown satisfactory retest reliability and the diffusion paradigm has been widely utilized in a wide range of behavioral tasks assessing various cognitive processes, including attention, lexical decision, recognition memory, priming, and numerical cognition (10–13). Moreover, this robust diffusion framework has been widely applied in clinical and non-clinical domains, including schizophrenia, attention deficit/hyperactivity disorder (ADHD), visual impairments (8, 14–16), and in brain-based interventions to target specific mental processes (11). It has been suggested that diffusion models can facilitate the understanding of clinical disorders (17). Thus, the application of this model has spanned disparate fields and holds the promise of illuminating the elusive function and malfunction processes of the central nervous system.

While different types of drift–diffusion models exist, one of the models, the EZ diffusion model, generates three unobserved variables: mean drift rate (cognitive processing), boundary separation (decision processing), and non-decision time (visuomotor processing). The mean drift rate is typically computed in simple binary choice tasks where humans accumulate evidence in favor of the different alternatives before committing to a decision, reflecting cognitive processing (18). Mean drift rate is a computational construct that assesses quality of information processes or speed of information uptake processes, where an increase in mean drift rate is believed to induce more accurate and faster decisions (19). Consistent with this, prior research showed that the mean drift rate is dynamic across human development. Boundary separation is a computational construct that assesses response conservativeness regarding the decision criterion. The trade-off between decision speed and accuracy is thought to be created by changing the boundary separation (19). Non-decision time is a computational construct that assesses the subset of the reaction time devoted to perceptual and motor response execution. Namely, non-decision time is the time spent on processing other than the decision process and it is usually referred to as reflecting the early perceptual processing of the stimulus of interest and the implementation of the motor response once the decision process is completed, reflecting visuomotor processing (7, 20, 21). Similar to mean drift rate and boundary separation, non-decision time exhibits well-characterized developmental shifts. Children exhibited lower mean drift rate, higher boundary separation, and longer non-decision time compared to college-aged adults in numerical discrimination tasks (22, 23).

### Which brain regions are involved in the diffusion parameters?

Frontal and parietal regions are thought to track evidence accumulation before reaching a decision in a task-independent way (18, 22, 24). Moreover, a study combining fMRI and the drift–diffusion model identified that frontoparietal areas corresponded to the decision variables resulting from the downstream stimulus–criterion comparison independent of stimulus type (25).

The involvement of mainly frontal and parietal regions in encoding the diffusion parameters raises the possibility that neurodevelopmental changes within the frontoparietal network regions may shape the well-documented developmental fluctuations in diffusion parameters, especially in tasks that mainly recruit the frontoparietal network, such as attention and numerical processing. In a similar vein, a recent magnetic resonance imaging (MRS) study revealed that glutamate and GABA profiles within the classic frontoparietal network regions, namely, the intraparietal sulcus (IPS, see **Supplementary File 7B**) and the middle frontal gyrus (MFG, see **Supplementary File 7A**), explained current and predicted future developmental changes in numerical cognition (26). However, it is unclear how the frontoparietal neurochemical profile elucidates neurodevelopmental alterations in diffusion parameters.

Given the above well-established developmental fluctuations in the diffusion parameters over the life span, it has been proposed that developmental shifts in all three diffusion parameters are possible explanations for corresponding developmental shifts in cognitive performance (23). However, in a recent study using 18 different response time tasks, age differences in intelligence were accounted for by age differences in non-decision time (23). This raises the intriguing possibility that the link between the frontoparietal regions and cognition across development is at the visuomotor level (i.e., non-decision time), rather than the more cognitive level (i.e., mean drift rate). To assess these open questions, we utilized an attention network task (**Fig. 1, Task 1**) because (i) it relies mainly on the frontal and parietal regions, (ii) it is suited for the calculation of the three diffusion parameters, and (iii) its performance is sensitive to developmental changes (27). We focused primarily on glutamate and GABA as these neurotransmitters were shown to be involved in attention and attention-related pathologies, including ADHD, in both human and animal work ((28–33); for extended text and references, see **Material and Methods** section).

**Fig. 1.**
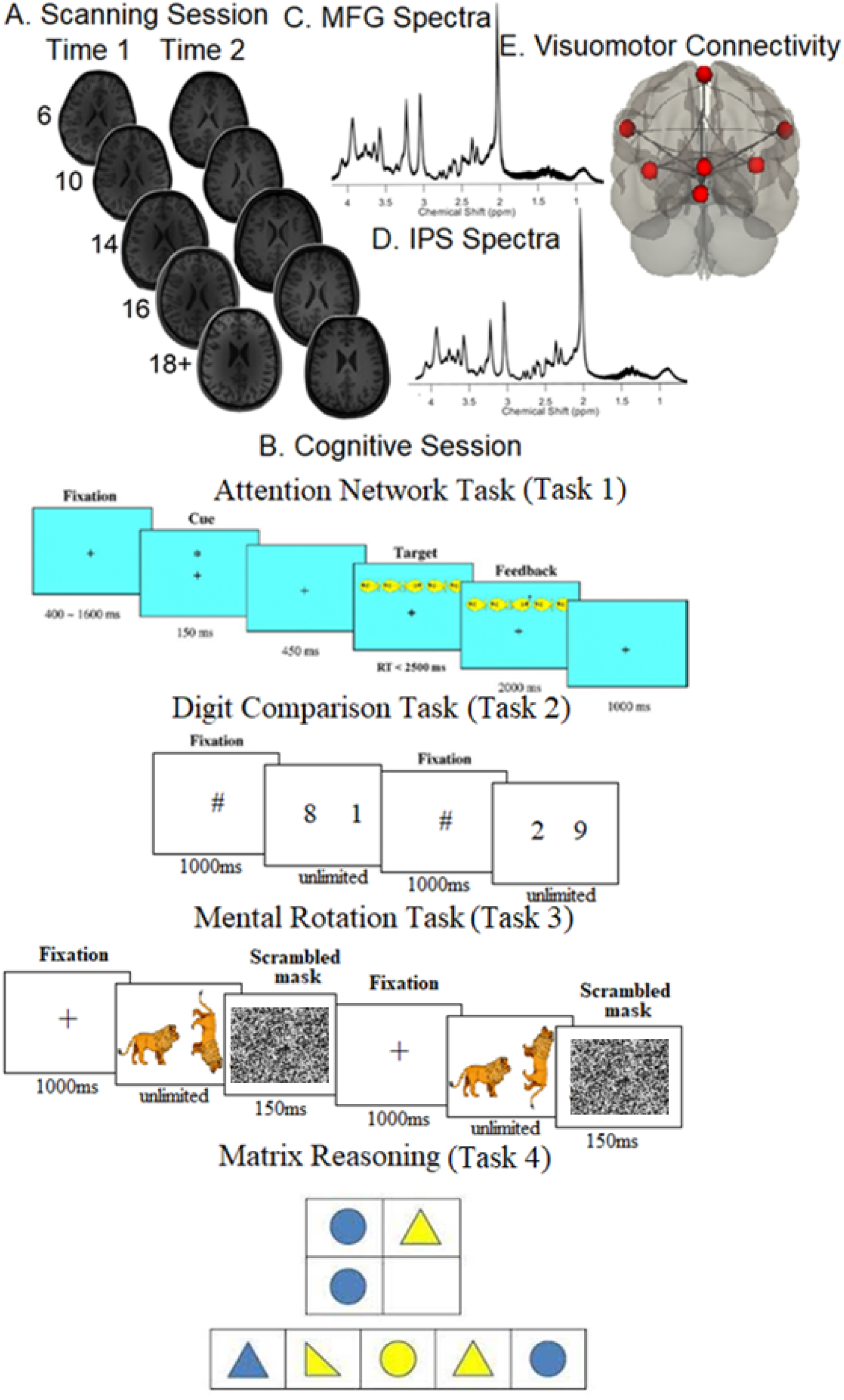
Scanning, cognitive session, neurochemical spectra plots, and visuomotor network connectivity. **A)** Scanning completed during the first assessment (A1) and the second assessment (A2, ~1.5 years later) in each of the five age groups (at A1: 6-year-olds, 10-year-olds, 14-year-olds, 16-year-olds, and 18+-year-olds). **B)** Cognitive session: *top panel*: the Attention Network task, where after a variable period, one of the possible cue types was presented for 150ms; after a pause of 450ms, one of the possible target types was presented, which required participants to indicate the direction of the middle fish arrow by pressing the corresponding key; *top middle panel*: the Digit Comparison task, where participants chose the larger value between two single-digit Arabic numbers. The numbers remained on the screen until the participant responded. Between each trial, a central fixation hashtag appeared for 1000ms. *Bottom middle panel*: the Mental Rotation task, where participants decided whether a rotated target figure presented on the right side of the screen was the same (i.e., non-mirrored) or different (i.e., mirrored) compared to the upright figure presented on the left side of the screen. Each trial began with a black fixation cross in the center of the white screen for 1000ms. *Bottom panel*: Matrix reasoning, which is a 30-item (of increasing difficulty) assessment tool that requires identifying a logical pattern in a sequence of visuospatial stimuli (this panel presents an example for illustration purposes only and does not report a trial from WASI II (91)). **C**-**D**) The mean spectra from our sample for the MFG (**C**) and IPS (**D**). The thickness corresponds to ±1 standard deviation from the mean (i.e., a chemical shift expressed in parts per million, ppm, on the x-axis). **E)** Brain regions consisting of the resting-state visuomotor functional connectivity network.

Given the critical role of the frontal and parietal regions, the role of development, and the role of neurochemicals in perceptual, cognitive, and motor tasks, several key research questions emerge: (i) How is frontoparietal neurochemical concentration related to mean drift rate, boundary separation, and non-decision time (henceforth, we will refer to non-decision time as visuomotor processing), and how do these associations change across human development? (ii) Do these associations between frontoparietal neurochemicals and visuomotor processing exist in a task-independent manner? In other words, do the associations between frontoparietal neurochemicals and visuomotor processing is replicated across different tasks such as numerical processing (**Fig. 1, Task 2**) and mental rotation (**Fig. 1, Task 3**)? (iii) Do the associations between frontoparietal neurochemicals and visuomotor processing explain individual differences in fluid intelligence?

A more optimal and informative way to address these questions is by using magnetic resonance spectroscopy (MRS). Several advantages of this technology make it suitable for studying these processes in the context of development. MRS has shown great promise in the classification and prediction of certain medical conditions in prior investigations. For example, it is known that neurochemical changes may precede anatomic changes (34). Therefore, it is possible to utilize and extend this unique predictive ability of MRS to discern which of these key neurobiological processes (cognitive, decision, visuomotor) are shaped by highly specialized neurochemical concentration across development. As mentioned above, our main aims were to examine whether neurochemical concentration within key frontoparietal regions can explain cognitive (mean drift rate), decision (boundary separation), and visuomotor processing. By testing participants from early childhood to early adulthood, we were able to examine whether these associations are static or dynamic across the lifespan.

## Results

### Task 1

#### 1.1. Developmental trajectories of the mean drift rate, boundary separation, and visuomotor processing

As a first step, we tracked the developmental trajectories of each of the three diffusion parameters (mean drift rate, boundary separation, visuomotor processing) during the first and the second assessment (for scatterplots, see **Supplementary File 2**; for statistical values, see **Supplementary File 4**). Lower scores in boundary separation and visuomotor processing indicate better performance, whereas higher scores in mean drift rate indicate better performance. As shown in **Supplementary File 4**, increasing age was associated with lower boundary separation and visuomotor processing and a higher mean drift rate, as expected. All these effects were replicated independently during the second assessment (**Supplementary File 4**).

#### 1.2. Neurochemicals and behavioral performance

After establishing the developmental trajectories of the three behavioral scores, we examined the capacity of neurochemicals to track these behavioral scores across development. For brevity, we present the neurochemical results that survived FDR correction and were neurochemically, anatomically, and diffusion-parameter specific (i.e., the results from assessing a given diffusion parameter hold even after controlling for the other two diffusion parameters) and replicated during the second assessment (see **Material and Methods** section for a complete description of this selection process).

There was a significant interaction between age and neurochemical measures in tracking visuomotor processing, which was replicated in the second assessment: IPS glutamate*age (first assessment: **Fig. 2A**, β=-.16, t(227)=-3.55, P_BO_=.007, CI=[−.28, −.04]; second assessment: **Fig. 2C**, β=-.24, t(154)=-3.47, P_BO_=.014, CI=[−.43, −.04]) and IPS GABA*age (first assessment: **Fig. 2B**, β=.09, t(227)=2.41, P_BO_=.033, CI=[.004, .17]; second assessment: **Fig. 2D**, β=.19, t(154)=3.29, P_BO_=.014, CI=[.06, .36]). Specifically, high glutamate predicted shorter visuomotor processing time in mature participants and longer visuomotor processing time in younger participants while GABA showed the opposite pattern.

**Fig. 2.**
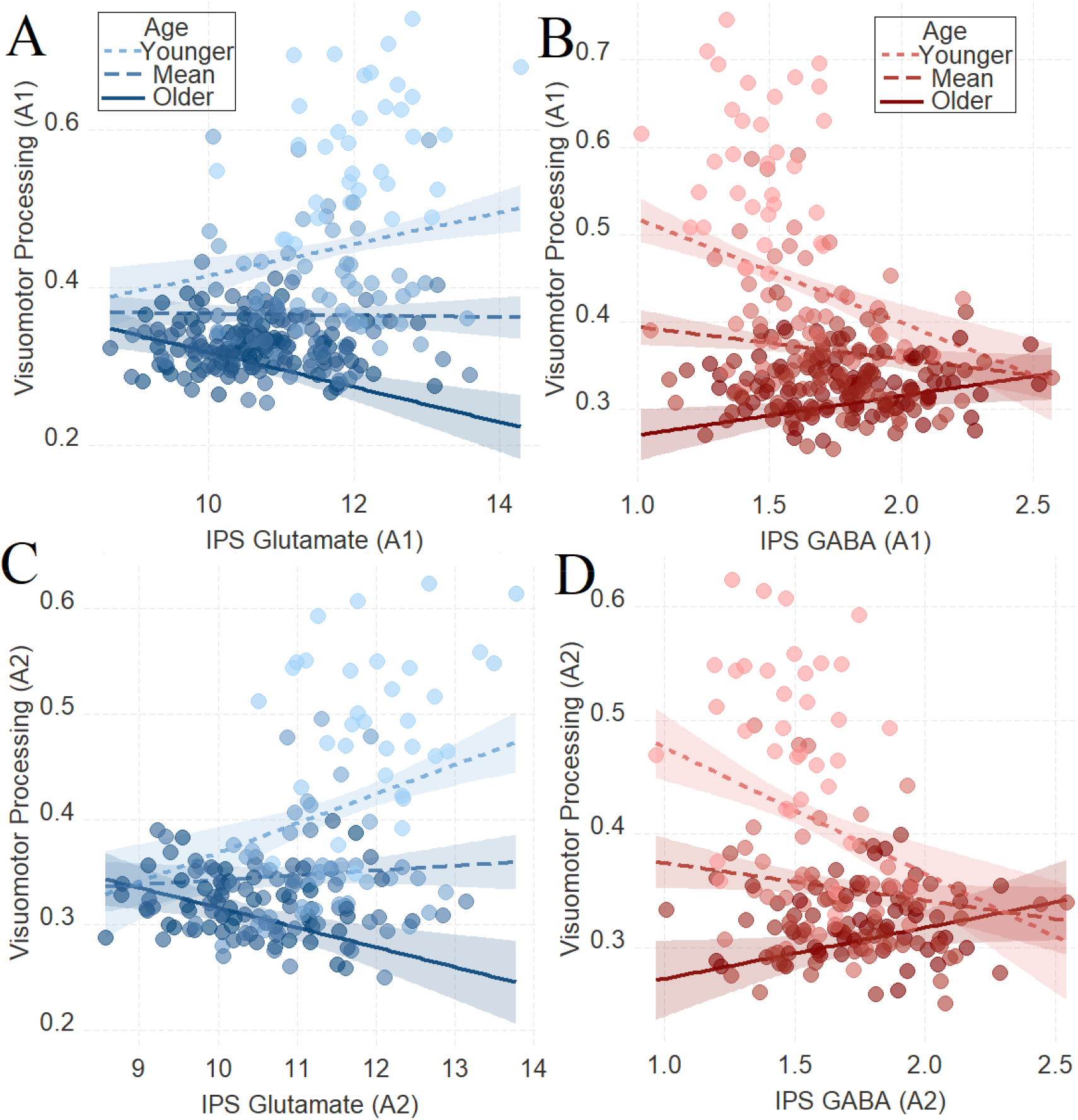
The moderating role of age in the relation between neurotransmitter concentration and visuomotor processing at A1 (A-B) and at A2 (C-D) in Task 1. To depict the interaction between the continuous variables (age and neurotransmitter concentration), we plotted the regression lines for younger and older participants (i.e., ± 1 standard deviation) from the mean age (92). Dark colors represent +1 SD above the mean (older participants), light colors represent −1 SD below the mean (younger participants), and medium colors represent the mean. **A**. IPS Glutamate*age and **B**. IPS GABA*age at A1; **C**. IPS Glutamate*age and **D**. IPS GABA*age at A2. For visualization purposes, we did not control for boundary separation and mean drift rate when plotting these panels.

### Task 2

The results of **Task 1** suggested that the neurochemical profile of individuals tracked individual variation in visuomotor processing across development and that these associations were neurochemically, neuroanatomically, and diffusion-parameter specific to visuomotor processing. The aim of **Task 2** was to replicate this set of findings that were obtained using the attention network task (**Task 1**) with a different task: a digit comparison task (see **Fig. 1B** and the **Materials and Methods section**).

#### 2.1. Developmental trajectories of the mean drift rate, boundary separation, and visuomotor processing

The behavioral results of **Task 2** mirrored the results of **Task 1**: increasing age was associated with lower boundary separation and visuomotor processing and with a higher mean drift rate (**Supplementary File 4.2**; for scatterplots, see **Supplementary File 2**).

#### 2.2. Neurochemicals and behavioral performance

As in **Task 1**, we found a significant interaction between age and neurochemical measures in tracking visuomotor processing in the task: IPS glutamate*age (first assessment: **Fig. 3A**, β=-.17, t(240)=-4.25, P_BO_<.001, CI=[−.26, −.09]; second assessment: **Fig. 3C**, β=-.16, t(170)=-3.17, P_BO_=.017, CI=[−.28, −.02]) and IPS GABA*age (first assessment: **Fig. 3B**, β=.18, t(240)=5.10, P_BO_<0.01 CI=[.08, .27]; second assessment: **Fig. 3D**, β=.29, t(168)=6.02, P_BO_<.001, CI=[.18, .4]).

**Fig. 3.**
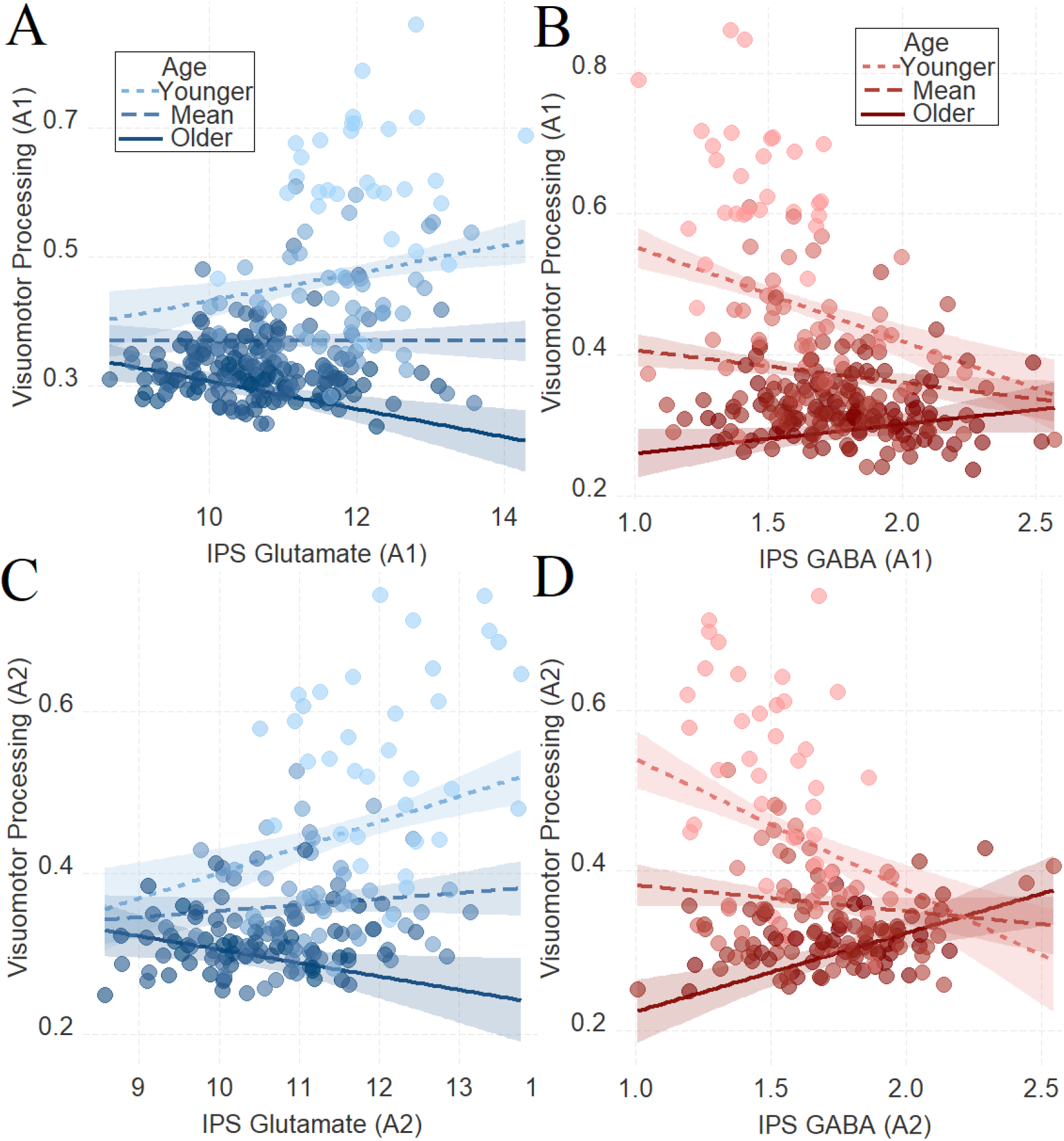
The moderating role of age in the relation between neurotransmitter concentration and visuomotor processing at A1 (A-B) and at A2 (C-D) in Task 2. To depict the interaction between the continuous variables (age and neurotransmitter concentration), we plotted the regression lines for younger and older participants (i.e., ± 1 standard deviation) from the mean age (92). Dark colors represent +1 SD above the mean (older participants), light colors represent −1 SD below the mean (younger participants), and medium colors represent the mean. **A**. IPS Glutamate*age and **B**. IPS GABA*age at A1; **C**. IPS Glutamate*age and **D**. IPS GABA*age at A2. For visualization purposes, we did not control for boundary separation and mean drift rate when plotting these panels.

### Task 3

The results of **Tasks 1–2** combined suggested that the neurochemical profile of individuals tracked individual variation in visuomotor processing across development. In **Task 3** we further examined this set of findings in a yet different task: a mental rotation task (see **Fig. 1B** and the **Materials and Methods section**).

#### 3.1. Developmental trajectories of the mean drift rate, boundary separation, and visuomotor processing

The behavioral results of **Task 3** reflected the results of **Tasks 1–2** in that increasing age was associated with lower boundary separation and visuomotor processing and a higher mean drift rate (**Supplementary File 4.3**; for scatterplots, see **Supplementary File 2).**

#### 3.2. Neurochemicals and behavioral performance

Similar to **Tasks 1–2**, we found a significant interaction between age and neurochemical measures in tracking visuomotor processing: IPS glutamate*age (first assessment: **Fig. 4A**, β=-.27, t(222)=-5.02, P_BO_<.001, CI=[−.42, −.11], which was not significant in the second assessment: **Fig. 4C**, β=.11, t(164)=1.67, P_BO_=.21, CI=[−.05, .3]) and IPS GABA*age (first assessment: **Fig. 4B**, β=.27, t(224)=4.66, P_BO_=001, CI=[.11, .43]; second assessment: **Fig. 4D**, β=.39, t(166)=5.46, P_BO_<.005, CI=[.22, .56]).

**Fig. 4.**
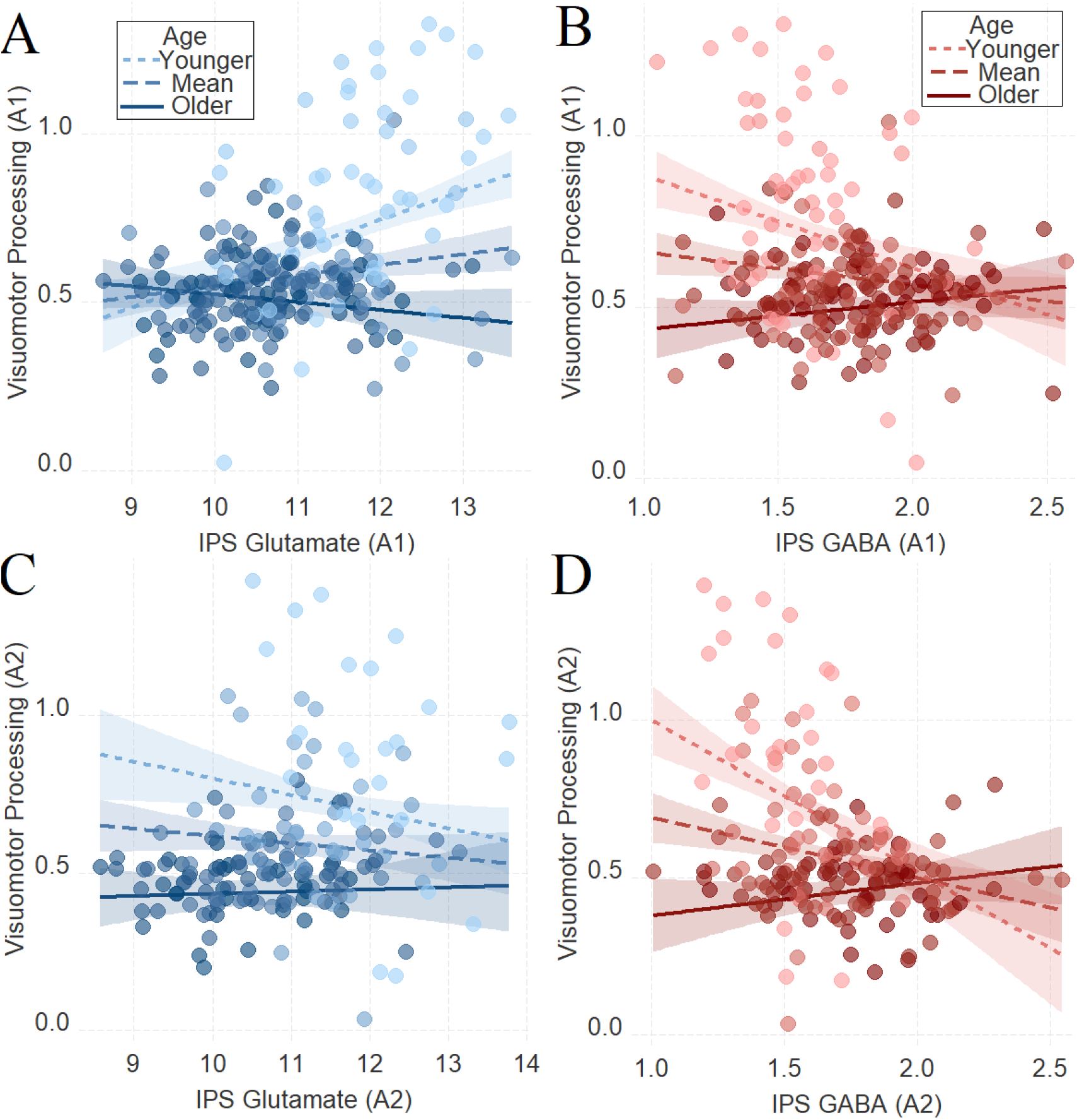
The moderating role of age in the relation between neurotransmitter concentration and visuomotor processing at A1 (A-B) and at A2 (C-D) in Task 3. To depict the interaction between the continuous variables (age and neurotransmitter concentration), we plotted the regression lines for younger and older participants (i.e., ± 1 standard deviation) from the mean age (92). Dark colors represent +1 SD above the mean (older participants), light colors represent −1 SD below the mean (younger participants), and medium colors represent the mean. **A**. IPS Glutamate*age and **B**. IPS GABA*age at A1; **C**. IPS Glutamate*age and **D**. IPS GABA*age at A2. For visualization purposes, we did not control for boundary separation and mean drift rate when plotting these panels.

### 4. Neurochemicals, visuomotor connectivity, and behavioral performance

After establishing the impact of IPS glutamate and GABA on tracking visuomotor processing in the same way across the three tasks as a function of development, we examined whether this relationship is influenced by visuomotor connectivity (see the **Materials and Methods** section for further details on the composite visuomotor processing score, which reflects visuomotor processing across the three tasks). We found that visuomotor connectivity interacted with age in tracking visuomotor processing (**Fig. 5C**, β=-0.16, T(215)=-3.9, SE=0.05, P_BO_=.002, CI=[−0.26, −0.06]) and we also found that IPS glutamate (**Fig. 5A**, β=−0.18, T(242)=-2.64, SE=0.08, P_BO_=0.022, CI=[−0.33, −0.02]) and IPS GABA (**Fig. 5B**, β=0.15, T(243)=2.10, SE=0.06, P_BO_=0.017, CI=[0.02, 0.26]) interacted with age in tracking visuomotor connectivity.

**Fig. 5.**
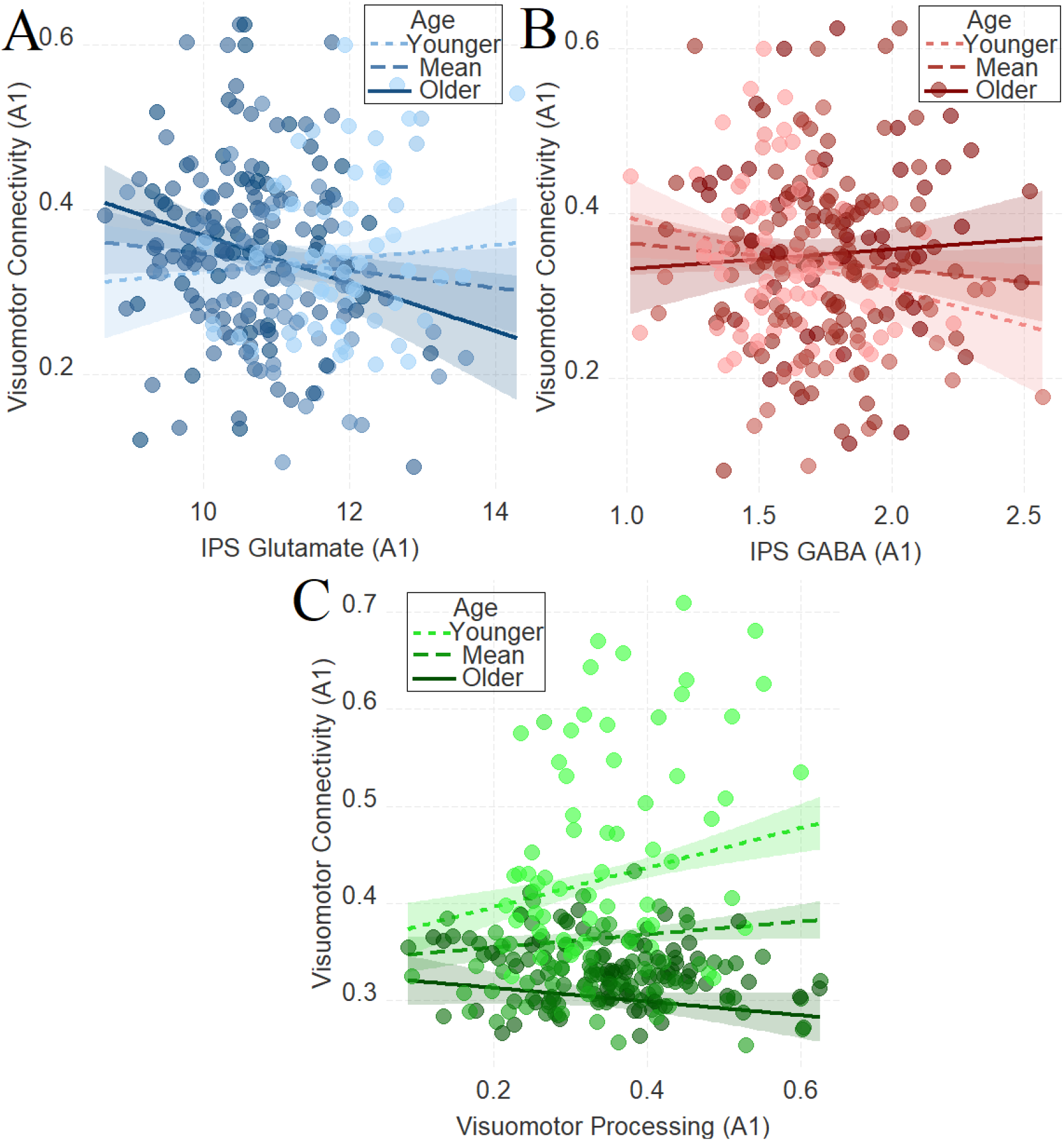
The moderating role of age in the relation between neurochemical concentration and visuomotor connectivity in IPS Glutamate (A) and GABA (B), and the moderating role of age in the relation between visuomotor connectivity and visuomotor processing (C). To depict the interaction between the continuous variables (age and neurotransmitter concentration), we plotted the regression lines for younger and older participants (i.e., ± 1 standard deviation) from the mean age (92). Dark colors represent +1 SD above the mean (older participants), light colors represent −1 SD below the mean (younger participants), and medium colors represent the mean. **A**. IPS Glutamate*age and **B**. IPS GABA*age at A1; **C**. IPS Glutamate*age and **D**. IPS GABA*age at A2. For visualization purposes, we did not control for boundary separation and mean drift rate when plotting these panels.

This set of findings raises the possibility that IPS neurochemicals (i.e., glutamate and/or GABA) shape visuomotor processing via visuomotor connectivity across development. To assess this possibility, we ran a moderated mediation (model 59; see **Material and Methods** section for details) where we evaluated whether the neurotransmitter concentration (independent variable) tracks visuomotor processing (dependent variable) via the visuomotor connectivity (mediator) as a function of development (moderator).

We found that the involvement of GABA and glutamate depends on the developmental stage: the relationship between IPS GABA and visuomotor processing is mediated by visuomotor connectivity only for the younger participants (indirect=−.03, CI=[−.06, −.001]; for full details, see **Supplementary File 8.1**). By contrast, visuomotor connectivity showed a trend in mediating the relationship between IPS glutamate and visuomotor processing only for the older participants (indirect=.02, CI=[−.003, .06]; for full details see **Supplementary File 8.2**).

### 5. Neurochemicals and intelligence scores

As mentioned in the Introduction, a behavioral study found that age differences in intelligence were accounted for by age differences in visuomotor processing (23). Importantly, the IPS has been highlighted as one of the brain hubs for fluid intelligence (35). Since we identified a strong and task-independent developmental effect between IPS neurochemicals and visuomotor processing, we utilized **Task 4** to assess whether visuomotor processing mediates the relationship between neurochemicals and fluid intelligence (see **Fig. 1B** and the **Materials and Methods section**).

As in **Tasks 1–3**, there was a significant interaction between age and glutamate in tracking intelligence scores: IPS glutamate*age (first assessment: **Fig. 6A**, β=.16, t(256)=3.72, P_BO_<.001, CI=[.06, .25]; second assessment: **Fig. 6C**, β=.12, t(178)=2.09, P_BO_=.02, CI=[.02, .23]). There was a significant interaction between age and GABA in tracking intelligence scores during the second assessment: IPS GABA*age (first assessment: **Fig. 6B**, β=-.05, t(257)=-1.09, P_BO_=.27, CI=[−.13, .04]; second assessment: **Fig. 6D**, β=-.15, t(178)=-2.40, P_BO_=.01, CI=[−.25, −.05]).

**Fig. 6.**
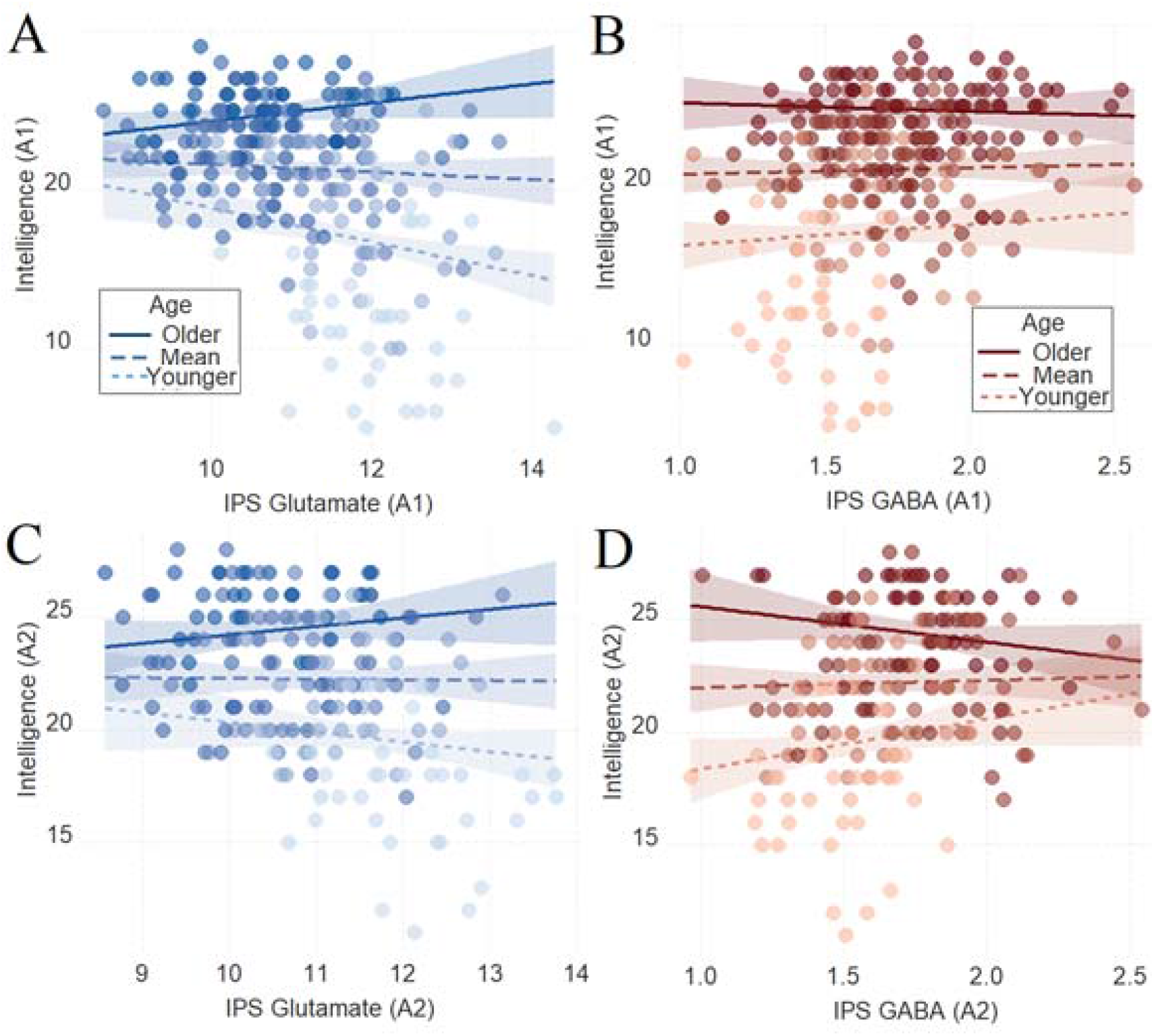
The moderating role of age in the relation between neurotransmitter concentration and intelligence at A1 (A-B), and at A2 (C-D) in Task 4. To depict the interaction between the continuous variables (age and neurotransmitter concentration), we plotted the regression lines for younger and older participants (i.e., ± 1 standard deviation) from the mean age (92). Dark colors represent +1 SD above the mean (older participants), light colors represent −1 SD below the mean (younger participants), and medium colors represent the mean. **A**. IPS Glutamate*age and **B**. IPS GABA*age at A1; **C**. IPS Glutamate*age and **D**. IPS GABA*age at A2. For visualization purposes, we did not control for boundary separation and mean drift rate when plotting these panels.

This set of findings raises the possibility that IPS neurochemicals shape fluid intelligence via visuomotor processing across development. To assess this possibility, we ran a moderated mediation (model 59; see **Material and Methods** section for details) where we evaluated whether the neurotransmitter concentration tracking of fluid intelligence as a function of age is mediated by visuomotor processing.

Visuomotor processing mediated the relationship between IPS GABA and fluid intelligence only for the older participants (**Fig. 7A**, indirect=.04, CI=[.002, .1]; **Fig.** for full details see **Supplementary File 8.3**). Similarly, visuomotor processing mediated the relationship between IPS glutamate and fluid intelligence only for the older participants (**Fig. 7B**, indirect=−.06, CI=[−.12, −.006]; **Fig.** for full details see **Supplementary File 8.4**).

**Fig. 7.**
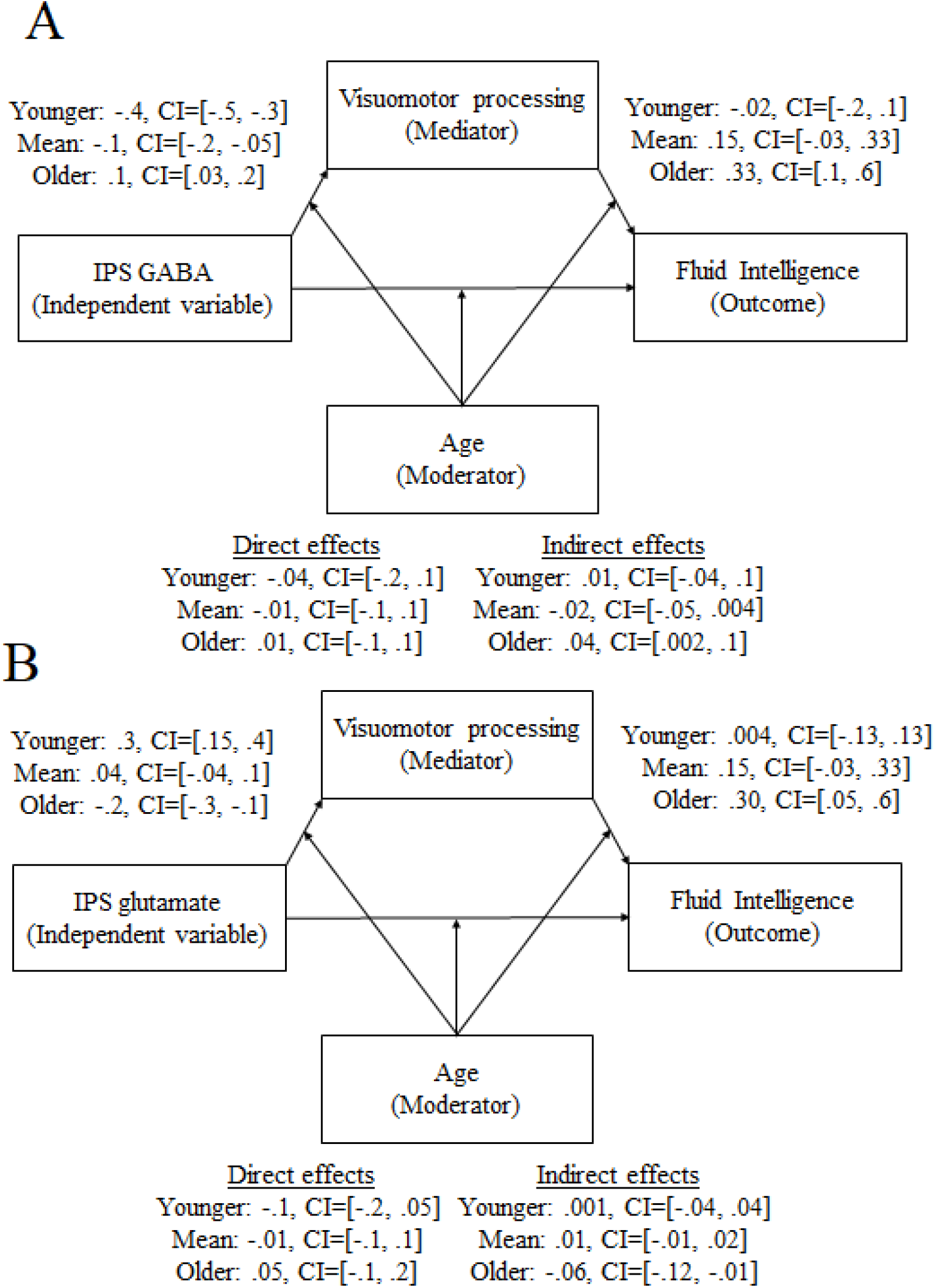
The moderating/mediating role of visuomotor processing in the relation between fluid intelligence and IPS GABA (A) or IPS glutamate (B) only for the older participants.

## Discussion

The present study examined the neurochemical mechanisms underlying cognitive, decision, and visuomotor processing across development by focusing on glutamate and GABA within the frontoparietal network. Four main findings emerged from our study: (i) visuomotor processing, as assessed with the computational metric non-decision time, is associated with glutamate and GABA within the IPS in a developmentally dependent manner, and was replicated in two additional tasks involving non-identical cognitive processing; (ii) the link between visuomotor resting-state connectivity and visuomotor processing, as well as IPS glutamate and GABA, is developmentally dependent; (iii) visuomotor connectivity mediates the relationship between IPS neurochemicals and visuomotor processing; and (iv) visuomotor processing mediates the relationship between IPS neurochemicals and fluid intelligence depending on the developmental stage.

### Visuomotor processing is associated with IPS glutamate and GABA in a developmentally dependent manner

Our results reveal how IPS glutamate and GABA, the neurotransmitters involved in brain excitation and inhibition (36–39), are associated with visuomotor functions in a developmentally dependent manner. Across three cognitive tasks, we showed that high glutamate predicted shorter visuomotor processing time in mature participants and longer visuomotor processing time in younger participants, while GABA showed the opposite pattern. These findings suggest that the relationship between IPS glutamate and GABA and visuomotor processing across development is task-independent.

Previous studies have examined the role of developmentally dependent glutamate levels in cognitive, emotional, and, importantly, motor functions (40–48). However, these studies focused primarily on adulthood and suggested that alterations in the availability of biochemical glutamate might contribute to the neural mechanisms underlying age-related cognitive and motor impairments (49). Our findings that in later developmental stages, higher levels of IPS glutamate are associated with better visuomotor performance (**Fig. 2A**, **2C**, **Fig. 3A**, **3C**, **Fig. 4A**, **4C**) are overall in line with the previous studies in older adults and extend them (49). Importantly, we obtained a negative relationship in the early developmental stages, where higher levels of IPS glutamate were associated with poorer visuomotor performance (**Fig. 2A**, **2C**, **Fig. 3A**, **3C**, **Fig. 4A**, **4C**). A possible explanation, as we also mention below, is that these glutamate-induced visuomotor transformations within the human IPS from early childhood to early adulthood may specifically occur within the IPS neurons, identified both in the human and in the non-human primate brain, that respond to the visuomotor domain (50). The accuracy of this working hypothesis can be tested in future animal pharmacological studies capable of reaching the necessary neuronal resolution that was beyond the reach of the present study using analogous tasks.

Regarding GABA, the maturation of GABA circuits and in particular the maturation of parvalbumin cells, a positive subtype of GABA neurons, is thought to be one of the molecular signatures triggering the onset of sensitive periods and plasticity, where an experimental increase or reduction of GABA triggers a precocious or delayed onset of sensitive periods/plasticity respectively (26, 51). One possibility, which we had suggested previously in the context of high-level cognition and we extended here to visuomotor processing (26), is that elevated GABA in early development may indicate greater plasticity, leading both to better visuomotor processing (i.e., shorter non-decision time) and to less positive connectivity in the visuomotor network. Regarding the relation between GABA levels and visuomotor processing in mature individuals, our findings indicate the opposite pattern of associations: elevated GABA leads both to poorer visuomotor processing (i.e., longer non-decision time) and to more positive connectivity in the visuomotor network.

Previous studies on GABA and cognition in adults, using modest sample sizes, yielded some conflicting results in that some studies found reduced GABA levels to be associated with cognitive improvement while others found the opposite pattern. For example, reduced GABA was associated with learning improvement in the motor system (52–54) and the visual system (55, 56), although some types of visual learning, and sensory learning in the tactile system, were associated with increased GABA (56–59). However, there are several reasons for these apparent discrepancies. For example, it is important to mention that the target MRS brain regions vary between these studies, and one cortical area may not represent or generalize to other cortical areas in terms of the association of neurotransmitter levels with the process under investigation. Besides these discrepancies, our findings in the mature participants suggest that lower GABA concentration within the IPS is associated with enhanced visuomotor processing, thus extending the involvement of GABA in the acquisition of visuomotor abilities. In addition to our suggestion of the involvement of developmental GABA in visuomotor processing, administration of GABAB agonist baclofen was shown to impair visuomotor processing in healthy adults (60). Consistent with this, we showed here that elevated GABA levels in the IPS are associated with poor visuomotor processing in later developmental stages. Importantly, because we employed a cross-sectional design, we were able to show that the relationship between GABA and visuomotor processing changes between early childhood and early adulthood. Although our sample included healthy individuals with a varied range of abilities, this finding can have implications for methods and applications aimed at detecting developmental atypicalities, and at altering GABA levels in earlier developmental stages in the case of visuomotor disorders.

The obtained results highlight the involvement of the IPS, but not the MFG, in visuomotor processing. These findings are in line with previous studies that have shown that visuomotor processing in the macaque and human IPS supports goal-directed behavior in a modality-general manner (61–64). Our findings support and extend this observation by identifying specific neurobiological markers within the IPS (i.e., glutamate and GABA) that are associated with goal-directed behavior from childhood to adulthood across four tasks. Moreover, our findings extend the results of previous studies, which were conducted using several modalities (39, 52–54, 60, 65–67), by showing that the involvement of GABA in visuomotor processing extends beyond the sensorimotor cortex and into the IPS.

### Visuomotor connectivity mediates the relationship between IPS neurochemicals and visuomotor processing

To delve deeper into the mechanistic network level, we employed resting fMRI and calculated the within-network connectivity of the visuomotor network for each participant. This visuomotor connectivity was associated with visuomotor performance and with IPS neurochemicals, providing a novel mechanistic account by which IPS neurochemicals, visuomotor networks, and development may be used to track the visuomotor functions of the human brain. Our findings support previous accounts suggesting that IPS serves as an interface between the sensory and motor systems (61), thereby revealing a developmentally dynamic and dissociable function of glutamate and GABA in shaping visuomotor processing that is replicable in multiple cognitive tasks.

The multimodal mediation analyses revealed that IPS glutamate and IPS GABA track visuomotor processing, as reflected by non-decision time scores, via visuomotor connectivity in a developmentally dependent manner that builds on previous studies. In particular, several previous studies reported animal and human evidence on the role of IPS in visuomotor functions (61, 68–70). Importantly, the human IPS was shown to subserve visuomotor transformation independently of the modality-specific processing of visual or proprioceptive information (61). Here we propose a generic, resting-state mechanistic model by which the levels of excitation and inhibition within the left IPS shape task-independent visuomotor functions through the visuomotor resting-state connectivity depending on the individual’s developmental stage. Specifically, our analysis revealed that visuomotor connectivity mediated the relationship between IPS GABA and visuomotor processing only for the younger participants, and visuomotor connectivity showed a trend in mediating the relationship between IPS glutamate and visuomotor processing only for the older participants. However, several aspects of this proposed model are still not known. For example, (i) are there specific time windows during visuomotor processing when the levels of IPS excitation and inhibition regulate the activity of visuomotor regions? Do the levels of IPS excitation and inhibition regulate the activity of visuomotor regions by regulating local GABA and glutamate within visuomotor regions? Addressing these questions is beyond the scope of the present study but can be examined in an investigation combining functional MRS and task-based fMRI (71, 72).

### Visuomotor processing mediates the relationship between IPS neurochemicals and fluid intelligence depending on the developmental stage

In a previous study, age differences in intelligence were accounted for by age differences in visuomotor processing (23). This raises the intriguing possibility that the developmental link we found between the IPS glutamate and GABA and fluid intelligence may be mediated by visuomotor processing. Indeed, we found that individual variation in visuomotor processing mediated the relationship between IPS glutamate and GABA levels and fluid intelligence in a developmentally dependent manner. Specifically, visuomotor processing mediated the relationship between IPS glutamate and GABA and fluid intelligence only for the older participants. Therefore, our study provides the neurobiological root that may drive this association: IPS glutamate and GABA. It is currently an open question why this relationship was observed only for the older participants. One potential explanation is that visuomotor processing becomes relevant and mediates the relationship between IPS neurochemicals and fluid intelligence only when the individual’s visuomotor processing has matured to the point that it becomes relatively automated and stabilized, which is the case with the older participants in our sample. Regarding the underlying neurobiological mechanisms, visuomotor processing becomes relevant in tracking fluid intelligence only when the visuomotor functions of the IPS have reached relative maturity. Specifically, the GABA- and glutamate-dependent maturity of IPS neurons that respond to the visuomotor domain may determine the onset of the influence of visuomotor processing on fluid intelligence (50). An alternative suggestion is that the strategies used to solve our fluid intelligence task differ as a function of development and are subserved by different mechanisms.

### Quantitative and qualitative differences in how participants solve fluid intelligence problems

It is noteworthy that even though the other two diffusion parameters (mean drift rate and boundary separation) are not related to IPS neurochemicals, the relationship between the three diffusion parameters and fluid intelligence was moderated by age (see **Supplementary File 9**). In particular, mean drift rate explained fluid intelligence only for the younger and the mean age groups, decision boundary explained fluid intelligence only for the younger age group, and non-decision time explained fluid intelligence only for the older age group. Moreover, the IPS is involved in fluid intelligence and visuomotor processing in later developmental stages (i.e., for the older but not the younger participants in our sample). Taken together, our results show that some of the mechanisms in fluid intelligence are based on cognitive (mean drift rate), decision (boundary separation), and visuomotor processing (non-decision time), and the involvement of each of these mechanisms depends on age. These findings suggest qualitative and quantitative differences in the way that participants across development solve fluid intelligence problems of the kind presented in our research.

In sum, we have shown that glutamate and GABA within the IPS track visuomotor functions across development. Using additional analyses we have established that visuomotor functional connectivity mediates the relationship between IPS neurochemicals and visuomotor performance, and that visuomotor performance mediates the relationship between IPS glutamate and GABA and fluid intelligence. These findings provide a mechanistic understanding of the involvement of GABA and glutamate in the IPS, and their potential contribution to other cognitive functions, such as mathematical development (26), which has been suggested to be subserved partly by visuomotor functions (73, 74). In addition, our study provides support for the view that fluid intelligence is also subserved by visuomotor processes, in this case in the older participants in our sample, and the involvement of glutamate and GABA. It is known that the brain develops and specializes to support such high-level cognitive functions (75–77). An integrative and robust understanding of psychology and biology across the different developmental stages, as we show in this study, would allow the researcher to progress toward personalized learning and precise diagnosis and intervention (78, 79).

## Materials and Methods

### Participants

We recruited 293 participants, and the demographic information for both the first assessment and the second assessment is reported in **Supplementary File 1**. The imaging session each lasted ~60min, and the cognitive and behavioral tasks described here each lasted ~30min and were part of a larger battery of tests. All imaging data were acquired in a single scanning session during which participants watched The LEGO Movie (80), except for the data acquired in the resting fMRI session (see below) during which participants were asked to fixate on a white cross displayed in the center of a black background. All participants were predominantly right-handed, as measured by the Edinburgh Handedness Inventory (81), and self-reported no current or past neurological, psychiatric, or learning disability or any other condition that might affect cognitive or brain functioning. For every assessment (i.e., first, second), adult participants received £50 compensation for their time, and children participants, depending on their age, received £25 (Early Childhood) and £35 (Late Childhood, Early Adolescence, Late Adolescence) in Amazon or iTunes vouchers, and additional compensation for their caregiver if the participant was under the age of 16. Informed written consent was obtained from the primary caregiver and informed written assent was obtained from participants under the age of 16, according to approved institutional guidelines. The study was approved by the University of Oxford’s Medical Sciences Interdivisional Research Ethics Committee (MS-IDREC-C2_2015_016).

### Magnetic Resonance Spectroscopy

Spectra were measured by semi-adiabatic localization using an adiabatic selective refocusing (semi-LASER) sequence (TE=32ms; TR=3.5s; 32 averages) (82, 83) and variable power RF pulses with optimized relaxation delays (VAPOR), water suppression, and outer volume saturation. Unsuppressed water spectra acquired from the same volume of interest were used to remove residual eddy current effects and to reconstruct the phased array spectra with MRspa (https://www.cmrr.umn.edu/downloads/mrspa/. Two 20×20×20mm^3^ voxels of interest were manually centered in the left intraparietal sulcus (IPS) and the middle frontal gyrus (MFG) based on the individual’s T1-weighted image while the participant was lying down in the MR scanner. Acquisition time per voxel was 10–15 minutes, including sequence planning and shimming.

MRS neurotransmitters were quantified with LCModel (84), using a basis set of simulated spectra generated based on previously reported chemical shifts and of coupling constants generated based on a VeSPA (versatile simulation, pulses, and analysis) simulation library (85). Simulations were performed using the same RF pulses and sequence timings as in the 3T system described above. Absolute neurochemical concentrations were extracted from the spectra using a water signal as an internal concentration reference.

The exclusion criteria for data were (i) Cramér–Rao bounds (CRLB) and (ii) the signal-to-noise balance (SNR). Neurochemicals quantified with Cramér–Rao lower bounds (CRLB, the estimated error of the neurochemical quantification) >50% were classified as not detectable. We aimed to follow a relatively unbiased approach, avoid using hard thresholds issues, and adopt the suggested procedure highlighted previously (86). In addition, since GABA concentration is relatively low (compared to glutamate), which usually induced high CRLB values, and since this study was mainly focused on GABA and glutamate, a good compromise was to exclude CRLB >50%. Additionally, we excluded cases with an SNR beyond 3 standard deviations per region. We also excluded cases with a neurochemical, connectivity (see below), or behavioral score beyond three standard deviations (per age group), and cases where the standardized residuals in a given analysis were beyond three standard deviations.

Absolute neurochemical concentrations were then scaled based on the structural properties of the selected regions and based on the predefined values shown in MRS-eq1 and MRS-eq2 (see below) (84); these predefined scaling values were therefore determined before the data collection. To quantify the structural properties, we segmented the images into different tissue classes including grey matter (GM), white matter (WM), and cerebrospinal fluid (CSF) using the SPM12 segmentation facility. Next, we calculated the number of GM, WM, and CSF voxels within the two masks of interest separately around the left IPS and MFG in native space. Subsequently, we divided these six numbers (GM, WM, and CSF for IPS and MFG, respectively) by the total number of GM, WM, and CSF voxels to obtain the corresponding GM, WM, and CSF fraction values per participant and region. As a final computation step, we scaled the absolute neurotransmitter values to these structural fractions using the following LCModel (84)) computation:

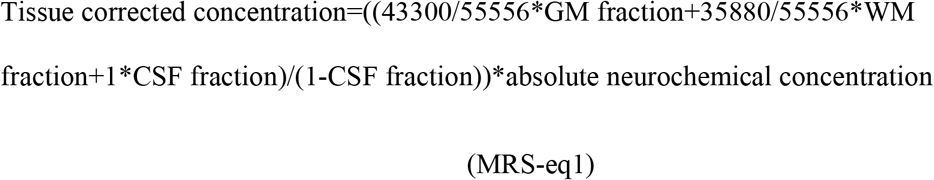

This was done to perform partial-volume correction based on three tissue classes as mentioned above. The numerator was the total water concentration in the region of interest and the factor 1/(1-CSF fraction) was the partial volume correction in that it corrected for the fact that all neurochemicals are concentrated in the grey and white matter. The values 43,300, 35,880, and 55,556 were the water concentrations in mmol/L for GM, WM, and CSF, respectively, and these were the default water concentrations values employed by the LCModel (87). The numerator corrected for differing tissue water concentrations for the unsuppressed water reference, whereas the denominator corrected for the assumption that CSF is free of metabolites.

These concentration values were scaled based on the T2 of tissue water values as can be seen in MRS-eq2 below. Fully relaxed unsuppressed water signals were acquired at TEs ranging from 32 to 4040ms (TR=15s) to water T2 values in each region of interest (32ms, 42ms, 52ms, 85ms, 100ms, 115ms, 150ms, 250ms, 450ms, 850ms, 1650ms, 3250ms, 4040ms). The transverse relaxation times (T2) of tissue water and the percent CSF contribution to the region of interest were obtained by fitting the integrals of the unsuppressed water spectra acquired in each region of interest at different TE values with a biexponential fit (88), with the T2 of CSF fixed at 740ms and three free parameters: T2 of tissue water, amplitude of tissue water, and amplitude of CSF water.

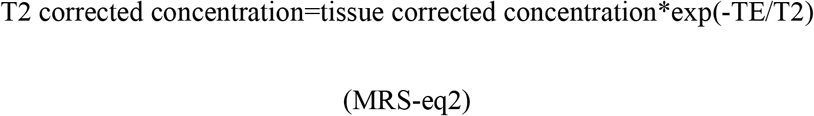

The results reported in the main text are derived using the quantification method of MRS-eq1. The results showed the same general pattern across all quantification methods.

### Resting functional magnetic resonance imaging (fMRI)

Functional images were acquired with a multi-band acquisition sequence (multi-band accel. factor=6, TR=933ms, TE=33.40ms, flip angle 64°, number of slices=72, voxel dimension=2×2×2mm, number of volumes=380). fMRI data were pre-processed and analyzed using the CONN toolbox (www.nitrc.org/projects/conn, RRID: SCR_009550) in SPM12 (Wellcome Department of Imaging Neuroscience, Institute of Neurology, London, UK) using the default pre-processing pipeline “MNI-space direct normalization” (89). Functional volumes were motion-corrected, slice-time corrected, segmented, normalized to a standardized (MNI) template, spatially smoothed with a Gaussian kernel (8mm FWHM), and pass filtered (0.01Hz to Inf). The low-pass portion of the filter (0.01Hz) was used to reduce the physiological and noise components contributing to the low-frequency segment of the BOLD signal, and the high-pass portion of the filter was set to Inf to additionally allow higher frequencies to be included (above .1Hz) given our fast acquisition. This also increased the degrees of freedom, thereby improving the quality of denoising and connectivity estimation. We also accounted for physiological noise in the time series by regressing out the confounding effects of white matter, CSF, realignment, and scrubbing that were automatically calculated in CONN. We calculated the functional connectivity of the visuomotor network [which is featured in CONN (89)], which consisted of seven regions, four of which were derived from the visual network [networks.Visual.Medial (2,-79,12), networks.Visual.Occipital (0,−93,−4), networks.Visual.Lateral (L) (−37,−79,10), networks.Visual.Lateral (R) (38,−72,13)], the three of which were derived from the motor network [networks.SensoriMotor.Lateral (L) (−55,−12,29), networks.SensoriMotor.Lateral (R) (56,−10,29), networks.SensoriMotor.Superior (0,-31,67)], yielding a single visuomotor functional connectivity score for each participant, which we refer to as visuomotor connectivity.

### Attention network task (Task 1)

Participants completed the child version of the original attention network task (90), and the stimuli were presented using E-Prime on a desktop computer (**Fig. 1B**). We included a single block of trials in the analyses (96 trials, 12 conditions), as this was the block that all age groups completed. We excluded trials with (i) a reaction time of <200ms, and/or with (ii) a reaction time that was beyond three standard deviations from the individual mean reaction time across all conditions. We calculated three behavioral measures (**EZ diffusion parameters**: mean drift rate, boundary separation, and visuomotor processing). The three EZ diffusion parameters were calculated following the procedure outlined in the original work (9) by averaging the mean reaction time of all correct trials without focusing on predefined conditions. For the results of the three attention network tasks separately, see the supplementary experiment in **Material and Methods** section.

### Digit comparison task (Task 2)

In the digit comparison task, participants were asked to choose the larger of two single-digit Arabic numbers as quickly and accurately as possible by clicking the mouse button corresponding to the side of the selected number (**Fig. 1B**). The numbers remained on the screen until the participant responded. Between each trial, a central fixation hashtag appeared for 1000ms. There were two practice trials followed by 72 test trials entailing all 36 possible comparisons of numbers between 1 and 9 repeated twice. The larger number appeared on the left side of the screen in half of the trials. For the results of the distance effect, see the supplementary experiment in **Material and Methods** section.

### Mental rotation task (Task 3)

In this task, participants decided whether a rotated target figure presented on the right side of the screen was the same (i.e., non-mirrored) or different (i.e., mirrored) compared to the upright figure presented on the left side of the screen (**Fig. 1B**). Each trial began with a black fixation cross in the center of the white screen for 1000ms. Thereafter, the two figures (upright and rotated) appeared on the screen, and participants responded by pressing the left or right mouse button, where the mouse button presses associated with the labels “same” and “different” were counterbalanced. The labels “same” and “different” appeared at the bottom left and bottom right of the screen, respectively, to remind participants which mouse button to press if the figures were the “same” or “different.” Participants were instructed to avoid mistakes and respond as fast as they could. The figures remained on the screen until response and then were followed by a scrambled mask for 150ms. There were two sets composed of 3 figures each (Animals set: giraffe, zebra, raccoon; Letters set: R, F, G). There were 24 trials for each figure, in which the rotated figure changed its pointing direction (left and right), degree of rotation (45, 90, 135), direction of rotation (clockwise and anticlockwise), and the rotated figure was either the same or different (non-mirrored or mirrored) compared to the upright figure. At the beginning of each block, participants completed ten practice trials, and they were allowed to take a break in the middle of each block. We focused on the “animal trials” as only these trials were present in all age groups. For the results of the distance effect, see the supplementary experiment in **Material and Methods** section.

### Fluid intelligence task (Task 4)

Participants completed the matrix reasoning subtest of the Wechsler Abbreviated Scale of Intelligence (WASI II) (91) as an index of fluid intelligence (**Fig. 1B**). The subset of matrix reasoning is a 30-item (of increasing difficulty) assessment tool that requires identifying a logical pattern in a sequence of visuospatial stimuli. The subtest is interrupted when the participant provides three consecutive wrong responses. We calculated the number of correct answers.

### Statistical Analyses

The dependent variables were three behavioural measures (i.e., the three **EZ diffusion parameters**: mean drift rate, boundary separation, and visuomotor processing). For the primary neurochemical analyses, the independent variables of interest were (i) the neurochemical concentration and (ii) the interaction between the neurochemical concentration and age. The former independent variable assesses whether there is an overall effect of neurochemical concentration on each diffusion parameter, while the latter independent variable considers the moderating role of age in shaping the relation between neurotransmitter concentration and that diffusion parameter. Importantly, in every model with an EZ diffusion parameter (e.g., mean drift rate) as the dependent variable, we controlled for the other two EZ diffusion parameters (e.g., boundary separation and visuomotor processing), as can be seen in eq1, eq2 and eq3 below, as this helped to assess the whether the results for one diffusion parameter were not confounded by the other two diffusion parameters. To assess these effects during the first assessment (A1) we employed eq1 (see below), to assess the same associations during the second assessment (A2) we employed eq2, and to assess the same associations during the prediction/longitudinal analyses we employed eq3. The prediction/longitudinal analyses were conducted to assess whether neurochemical concentration and its confluence with age during the first assessment can predict three behavioural measures (i.e., mean drift rate, boundary separation, and visuomotor processing) obtained during the second assessment. In the equations below, the “ behavioural score” is always one of the three EZ diffusion parameters, and the control behavioural score 1 and 2 are the other two EZ diffusion parameters.

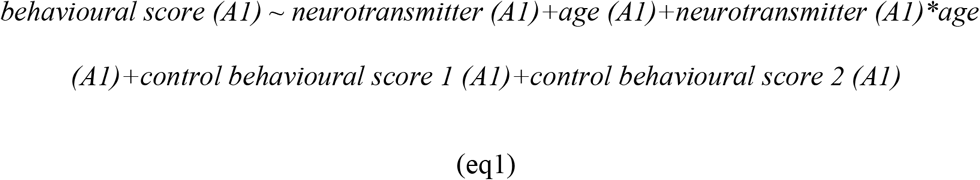

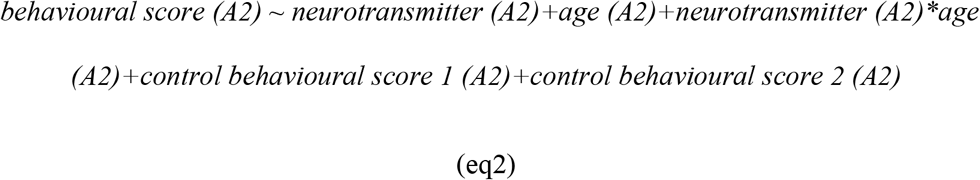

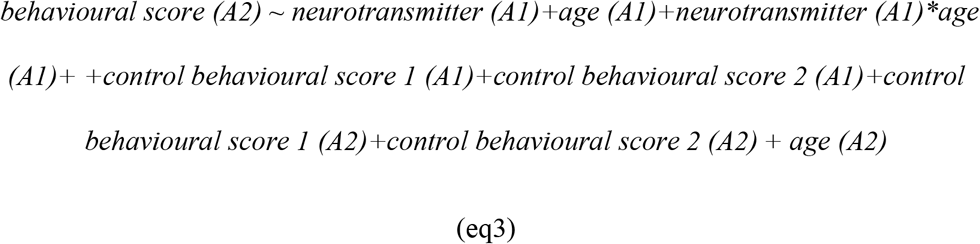

To correct for multiple comparisons during the first assessment, we performed FDR-correction correcting for 36 comparisons [**EZ diffusion parameter** (3: drift rate, boundary separation, visuomotor processing) \***neurochemical** (3: GABA, glutamate, NAA)\***region** (2: IPS, MFG)\***type of effect** (2: neurochemical*age interaction, the main effect of neurochemical)]. To correct for multiple comparisons during the prediction/longitudinal analyses, we performed FDR-correction correcting for 36 comparisons [**EZ diffusion parameter** (3: drift rate, boundary separation, visuomotor processing)\***neurochemical** (3: GABA, glutamate, NAA)\***region** (2: IPS, MFG)\***type of effect** (2: neurochemical*age interaction, the main effect of neurochemical)]. From the longitudinal analyses (**Task 1**), the only association that survived the FDR correction was the IPS GABA statistical model predicting visuomotor processing, but this did not survive the neurochemical and neurotransmitter specificity.

Since we applied a separate FDR multiple comparison correction for the first assessment analyses in **Task 1** and for the prediction/longitudinal analyses in **Task 1**, we did not correct for multiple comparisons for all replication analyses, nor did we assess the (diffusion parameter, neurochemical and neurotransmitter) specificity in the replication analyses of **Task 2** and **Task 3** to reduce type 2 errors. Regarding the supplementary experiment (for details, see **Material and Methods** section), we applied a separate FDR multiple correction correcting for 180 comparisons during the first assessment and a further separate FDR multiple correction correcting for 180 comparisons during the longitudinal/prediction analyses [**EZ diffusion parameter** (3: drift rate, boundary separation, visuomotor processing) \***neurochemical** (3: GABA, glutamate, NAA)\***region** (2: IPS, MFG)\***type of effect** (2: neurochemical*age interaction, the main effect of neurochemical)\***cognitive function** (5: alerting network, orienting network, executive network, distance effect, SNARC effect)].

Apart from correcting for multiple comparisons, we assessed the neurotransmitter and neuroanatomical specificity of our findings that survived the aforementioned FDR correction. To this end, we ran multiple regression models with several independent variables, namely the main effect of all six neurotransmitter measures (i.e., IPS glutamate, IPS GABA, IPS NAA, MFG glutamate, MFG GABA, MFG NAA) and their interactions with age, as well as the main effect of age as can be seen in eq4 (A1), eq5 (A2), and eq6 (prediction/longitudinal) below.

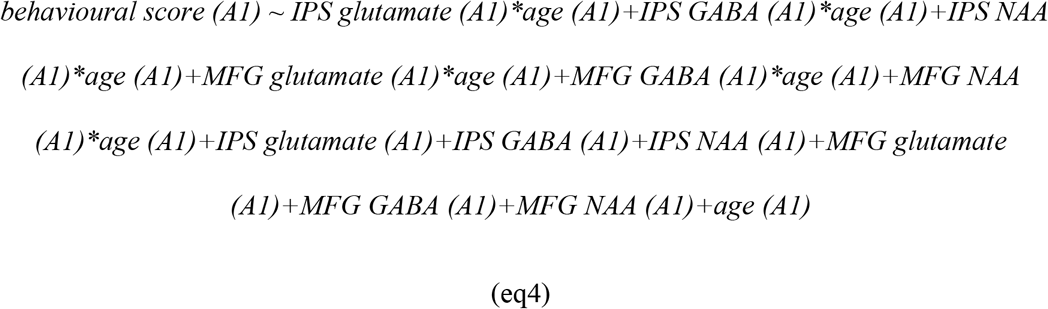

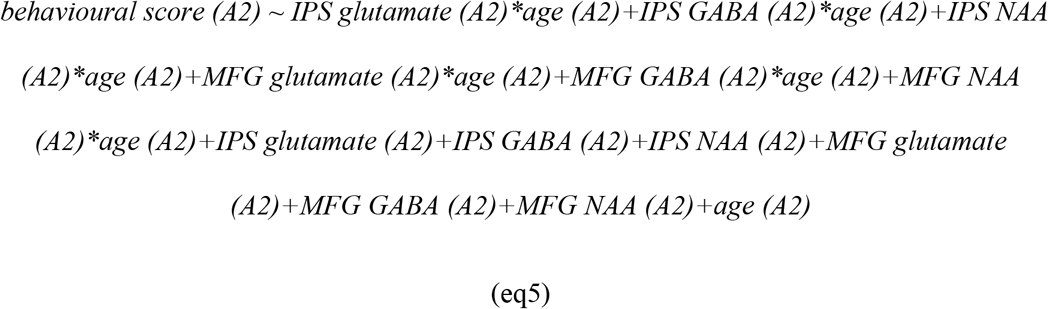

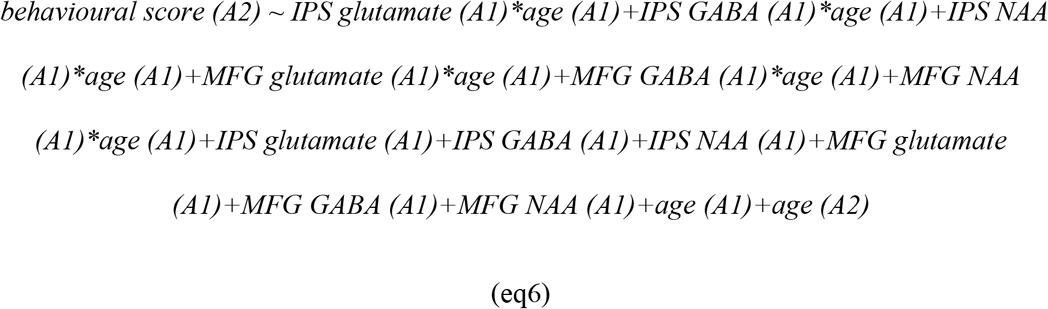

The statistical values in the main text were obtained from eq4, eq5, and eq6, but for the results from eq1 and eq2 see **Supplementary File 3**.

In **Task 1-3**, visuomotor processing score was calculated based on each task accordingly meaning visuomotor processing in **Task 1** was computed based on the attention network task (**Task 1**), visuomotor processing in **Task 2** was computed based on the digit comparison task (**Task 2**), and visuomotor processing in **Task 3** was computed based on the mental rotation task (**Task 3**). Since we obtained the same pattern of findings regarding visuomotor processing across **Task 1-3** which was one of our earlier aims was addressed, for the visuospatial connectivity analyses and all moderated mediation analyses, we computed a composite visuomotor processing score that was obtained based on **Task 1-3**. Specifically, we zscored the visuomotor processing in each of the three tasks separately and calculated the mean of these three scores which represents a generic or task-independent visuomotor processing score for each participant.

To assess whether the visuomotor connectivity interacted with age in explaining visuomotor processing, we run the multiple regression model displayed below in eq7.

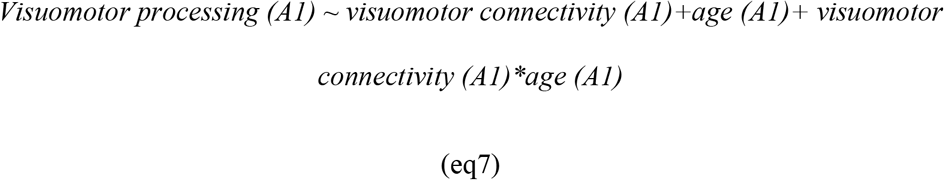

To assess whether the neurotransmitter scores interacted with age in explaining the visuomotor connectivity, we run a multiple regression model displayed below in eq8.

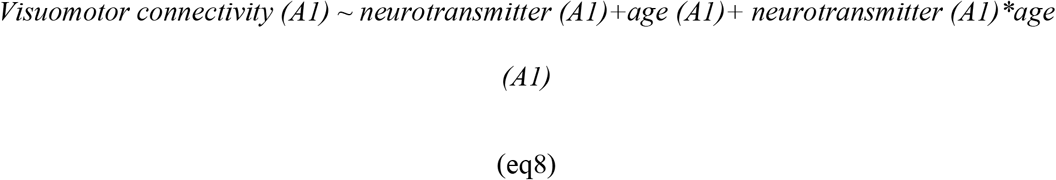

The statistical results presented in the results section involve multiple linear

regressions with bootstrapping (5000 samples) and show the uncorrected P-value denoted by P_BO_, as well as the lower (CI_L) and upper (CI_U) bound and bound of the confidence intervals obtained from bootstrapping.

To examine the possibility that IPS neurochemicals (i.e., glutamate and/or GABA) are associated with visuomotor processing via visuomotor connectivity across development, we ran a moderated mediation model (model 59, PROCESS v4.1, (30)) where we evaluated whether the neurotransmitter concentration (independent variable) tracks visuomotor processing (dependent variable) via the visuomotor connectivity (mediator) as a function of development (moderator).

In these mediation analyses and all other multiple regression models, we additionally removed cases that fell beyond three standard deviations after running the multiple regression models (in this case, eq9).

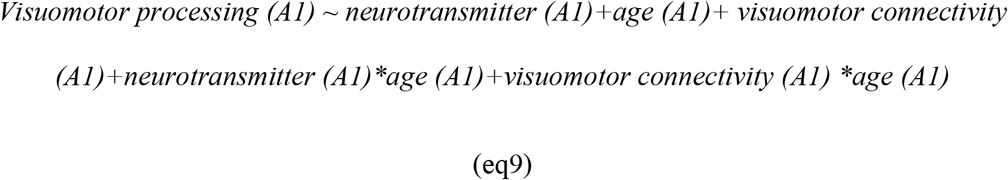

To examine the possibility that IPS neurochemicals were associated with intelligence via visuomotor processing across development, we ran a moderated mediation model (model 59, PROCESS v4.1, (30)) where we evaluated whether the neurotransmitter concentration (independent variable) tracks intelligence (dependent variable) via the visuomotor processing (mediator) as a function of development (moderator). We assessed the statistical significance by examining the indirect effect at 90% confidence intervals (and not all individual paths) with bootstrapping (5000 samples). The reason we used 90% was that our mediation analyses were follow-up analyses based on the significant results obtained in advance.

In these mediation analyses and all other multiple regression models, we additionally removed cases that fell beyond three standard deviations after running the multiple regression models (in this case, eq10).

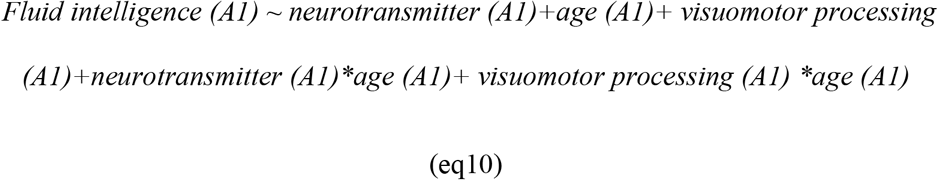

### Supplementary Experiment

Attention is vital for daily functioning and plays a critical role in human development across multiple domains, such as perception, language, and memory (1, 2). Altered attention processing manifests in severe disorders including, but not limited to, attention deficit hyperactivity disorder (ADHD), schizophrenia, borderline personality disorder, and brain injury (e.g., stroke) (3).

Several models have presented attention as a multifaceted construct typically divided into several functional systems (4). In particular, Posner’s model of attention entails three systems: (i) alerting, (ii) orienting, and (iii) executive (5, 6). The altering system is divided into two subcomponents: tonic and phasic (7). Tonic alerting corresponds to sustained attention or vigilance, which is needed to respond to low-frequency events rapidly [8]. On the other hand, phasic alertness corresponds to the advantage in processing stimuli preceded by warning cues compared with those that are not (6). The orienting system represents individuals ability to move attention to a spatial location (2). The executive system allows allocating attention to objects or events despite conflicting information and allows switching among tasks.

All three attention systems exist to some degree in infancy and change over lifespan (8). Cross-sectional studies have suggested that the executive system stabilizes around the age of 7, alertness improves even beyond the age of 10, whereas orienting remains stable (8). Other studies, examining children from 6 to 12 years, concluded that the alerting system matures early in development and the orienting and executive networks continue developing throughout childhood (9). Later in life, alerting decreased with age, whereas the other two systems remained stable in adulthood (10). Recent longitudinal studies revealed that alerting kept developing between the age of 8 and 10, while orienting stabilised at 6 and executive stabilized at 7 (11). Another longitudinal study in children with and without ADHD, who were assessed four times between the age of 7 and 11, revealed improvement in all attention systems apart from orienting. Moreover, typically developing children showed superior executive functioning compared to children with ADHD (12).

Recent evidence supports the existence of five brain networks underpinning the three attention systems, one brain network for the alerting system and two brain networks each for the orienting and the executive system (4). The locus coeruleus and its projections to the frontal and parietal regions underpin alerting (13, 14). The ventral attention network and the dorsal network, including the frontal eye fields and the intraparietal sulcus (IPS), underpin bottom-up orienting and top-down visuospatial functions, respectively (4, 15–18). The cingulo-opercular network and the frontoparietal network, mainly featuring the dorsolateral prefrontal cortex and the IPS, underpin the executive system (4).

Several pharmacological studies have been conducted to date to examine the three attention systems. Pharmacological manipulation has shown that blocking norepinephrine decrease alerting (19), whereas acetylcholine, via nicotine administration, reduced orienting performance (20). The administration of Adderall and Ritalin, which increase the concentration of dopamine and norepinephrine, improves executive control in individuals with ADHD. Animal work revealed the important role of glutamate and GABA, the brain’s major excitatory and inhibitory neurotransmitters, in attention (21).

As discussed in the Introduction, magnetic resonance spectroscopy (MRS) has emerged as a valuable tool to investigate the role of neurochemicals across development. MRS studies on attention mainly compared individuals with and without ADHD symptoms. Children (8-12 years) with ADHD exhibit reduced concentration of GABA in the somatosensory/motor cortex compared to typically developing children (22), possibly suggesting a deficit in short intracortical inhibition in ADHD. Glutamate was also investigated in the context of ADHD, where children with ADHD exhibited higher glutamate (23). Other MRS studies focused on N-Acetylaspartate (henceforth; NAA), a marker of neuronal integrity and health, yielding conflicting findings. For example, individuals with ADHD compared to controls, exhibit below-normal NAA levels in some studies and above-normal levels in other studies (24, 25). A meta-analysis identified a higher concentration of NAA in the prefrontal cortex in children with ADHD, but not in adults with ADHD, showing that the abnormal excess in NAA dropped linearly with increasing age, potentially resolving previous discrepancies between studies (26).

The assessment of the three attention systems is typically done using the Attention Network Task (5, 6) (**Fig 1**), which allows comparing reaction times in different conditions (see below) to obtain for each participant the altering, orienting, and executive network indexes. However, the use of these metrics prevents the researcher from discerning the underlying processes of attention. To put it simply, if 14-year-old children exhibit superior executive systems scores compared to 6-year-old children, the researcher cannot determine whether this difference depends on the cognitive, perceptual, motor, decision processing, or a combination of these from the single reaction time measure alone.

Apart from regulating the processing of the attention system, the frontoparietal network is also engaged when we compare quantities in the context of nonsocial comparisons (e.g., which of two numbers is larger in magnitude, which of two dogs is taller) as well as in the context of social comparison (e.g., which of two women is more attractive). A classic effect in the literature that applies in the context of the frontoparietal network is the distance effect, which states that the closer two compared magnitudes (e.g. two numbers), the more difficult the comparison, and the greater the activity of this frontoparietal network (27, 28). In the context of physical beauty, the distance effect was observed when comparing the beauty of two unknown people (29). In particular low-distance beauty comparisons elicited longer reaction times, lower accuracy, and greater recruitment of the frontoparietal network compared to high-distance beauty comparisons (29).

#### Aims of the supplementary experiment

(i) Given the crucial role of frontoparietal networks in regulating attention processing (see above) and for complementing the analyses of **Task 1**, we conducted a supplementary experiment that aimed to assess whether frontoparietal neurochemical concentration tracks and predicts the three EZ diffusion parameters (i.e., mean drift rate, boundary separation, and visuomotor processing; for details, including multiple comparison corrections, see **Material and Methods** section) within each of the three attention networks separately. The alertness network was calculated by subtracting the EZ diffusion parameter during the double-cue condition from the same EZ diffusion parameter in the no cue condition. The orienting network was calculated by subtracting the EZ diffusion parameter during the spatial cue condition from the same EZ diffusion parameter in the central cue condition. The executive network was calculated by subtracting the EZ diffusion parameter during the congruent condition from the same EZ diffusion parameter in the incongruent condition.

(ii) Given the crucial role of frontoparietal networks in regulating the distance effect (see above) and for complementing the analyses of **Task 2-3**, we examined whether frontoparietal neurochemical concentration tracks and predicts the three EZ diffusion parameters (i.e., mean drift rate, boundary separation, and visuomotor processing; for details, see **Material and Methods** section) in response to the distance effect [in **Task 2-3**, and for completeness in response to the spatial-numerical association of response codes (SNARC) effect in **Task 2**]. In **Task 2-3**, the distance effect was calculated by contrasting the high-distance vs low-distance trials (in **Task 2**: far trials vs near trials and in **Task 3**: 135 degrees trials vs 45 degrees trials).

Regarding the first assessment analyses of the supplementary experiment, the only association that survived the FDR correction in the supplementary experiment was the MFG NAA statistical model predicting decision boundary in response to the distance effect in **Task 2**, but this did not survive the neurochemical and neurotransmitter specificity. Regarding the longitudinal analyses of the supplementary, the only association that survived the FDR correction in the supplementary experiment was the IPS glutamate statistical model predicting decision visuomotor processing within the alerting network in **Task 1** (β=-0.30, T(143)=-2.55, SE=0.12, P_BO_=0.017, CI=[−0.54 −0.05]), and this result survived the neurochemical and neurotransmitter specificity.

For completeness, however, for the attention network results as well as for the distance and SNARC effects, see **Supplementary File 4** (behavioural age effects), **Supplementary File 5** (neurochemical effects without controlling for behavioural scores during the first assessment when applicable), and **Supplementary File 6** (neurochemical effects with controlling for behavioural scores during the first assessment when applicable).

## Acknowledgments

The authors are grateful to all the participants and caretakers involved in this study, and to Malin I. Karstens, Katarzyna Dabrowska, Laura Epton, Charlotte Hartwright, Ramona Kantschuster, Margherita Nulli, Claire Shuttleworth, and Anne Sokolich for their assistance in running this project. The authors are grateful to all Wellcome Centre for Integrative Neuroimaging (WIN) staff, in particular Nicola Filippini, Emily Hinson, Eniko Zsoldos, Caitlin O’Brien, Jon Campbell, Michael Sanders, Caroline Young, and David Parker. The Wellcome Centre for Integrative Neuroimaging (WIN) is supported by core funding from the Wellcome Trust (203139/Z/16/Z). This work was supported by the European Research Council (Learning&Achievement Grant 338065).

## Data availability

All quantitative data will be deposited in XNAT upon publication. A subset of the MRI data have been deposited in XNAT (https://central.xnat.org/data/projects/PN21).

